# Pathophysiology of hypereosinophilia-associated heart disease

**DOI:** 10.1101/2024.07.03.601845

**Authors:** Usman Sunusi, Ben Ziegelmeyer, Immaculeta Osuji, Mario Medvedovic, Haley Todd, Joe Abou-Khalil, Nives Zimmermann

## Abstract

**Background:** Cardiac complications in patients with hypereosinophilia cause significant morbidity and mortality. However, mechanisms of how eosinophilic inflammation causes heart damage are poorly understood.

**Methods:** We developed a model of hypereosinophilia-associated heart disease by challenging hypereosinophilic mice with peptide from the cardiac myosin heavy chain. Disease outcomes were measured by histology, immunohistochemistry, flow cytometry, and measurement of cells and biomarkers in peripheral blood. Eosinophil dependence was determined by using eosinophil-deficient mice (ΔdblGATA). Single cells from heart were subjected to single cell RNA sequencing to assess cell composition, subtypes and expression profiles.

**Results:** Mice challenged with myocarditic and control peptide had peripheral blood leukocytosis, but only those challenged with myocarditic peptide had heart inflammation. Heart tissue was infiltrated by eosinophil-rich inflammatory infiltrates associated with cardiomyocyte damage. Disease penetrance and severity were dependent on the presence of eosinophils. Single cell RNA sequencing showed enrichment of myeloid cells, T-cells and granulocytes (neutrophils and eosinophils) in the myocarditic mice. Macrophages were M2 skewed, and eosinophils had an activated phenotype. Gene enrichment analysis identified several pathways potentially involved in pathophysiology of disease.

**Conclusion:** Eosinophils are required for heart damage in hypereosinophilia-associated heart disease. Additionally, myeloid cells, granulocytes and T-cell cooperatively or independently participate in the pathogenesis of hypereosinophilia-associated heart disease.

## Introduction

Eosinophil-associated disorders (EADs), including hypereosinophilic syndrome (HES), eosinophilic granulomatosis with polyangiitis (EGPA) and eosinophilic gastrointestinal disorders (EGID), are a heterogeneous group of conditions characterized by blood and/or tissue hypereosinophilia and eosinophil-related clinical manifestations^1^. Cardiac complications occur in up to 60% of patients with sustained hypereosinophilia^2-4^, and are a major cause of morbidity and mortality in this patient population. In patients with eosinophilic heart disease (EHD), the clinical course is characterized by eosinophil-rich endomyocarditis with cardiomyocyte necrosis, followed by replacement fibrosis in the myocardium and possibly thrombosis stemming from endocardial damage, eventually leading to cardiomyopathy^3-7^.

As the NIH Taskforce on Research needs of Eosinophil-Associated Diseases (TREAD) and recent RE-TREAD reported, there is a paucity of preclinical models that adequately replicate cardiac disease in hypereosinophilia, and development of these models would enable mechanistic studies aiming to develop targeted therapies^8,9^. To address this unmet need, we have recently developed^10,11^ a mouse model of EHD that recapitulates many of the salient features of the human disease, importantly including hypereosinophilia with heart involvement reminiscent of that seen in patients. While informative, this model has several limitations including low and highly variable penetrance, unpredictable clinical course, and first presentation with sudden death. These limitations make mechanistic studies difficult. Diny et al^12^ challenged wild type and hypereosinophilic mice with cardiac myosin peptide, to induce experimental autoimmune myocarditis (EAM) and eosinophilic EAM (eoEAM). They showed that progression of myocarditis to dilated cardiomyopathy (DCM) is dependent on the presence of eosinophils, thus implicating them in the pathophysiology of disease. While this progression was dependent on eosinophil production of IL-4 in the EAM model (not associated with hypereosinophilia), the mechanism of eosinophil-mediated disease effects has not been studied in hypereosinophilic mice (eoEAM model), which showed significantly different levels of inflammation and cardiac dysfunction^12^. Notably, studies in eosinophil associated diseases (beyond heart disease) have shown that eosinophils may either contribute to tissue repair or tissue damage, which is likely disease dependent. Therefore, the focus of studies presented in this manuscript was to study the mechanism of eosinophil-mediated effects on heart function in hypereosinophilia, specifically which role eosinophils play.

In this manuscript we show that eosinophils are required for heart damage in hypereosinophilia-associated heart disease. Furthermore, we provide a detailed understanding of the cellular and molecular underpinnings of eosinophilic heart disease.

## Methods

### Mice

IL-5 transgenic mice (IL-5tg) in which the IL-5 gene is driven by the CD2 promoter^13^ on a BALB/c background were provided by Dr Marc Rothenberg (Cincinnati Children’s Hospital). Eosinophil-deficient ΔdblGATA mice^14^ (BALB/c background) were provided by Dr Rothenberg, with approval from Dr. Orkin. Mice were housed in specific pathogen-free facility at the University of Cincinnati. Experiments were conducted on >6wk-old mice of both genders; initial studies did not identify gender-specific differences in measured outcomes. All procedures and protocols were approved by the Institutional Animal Care and Use Committee of the University of Cincinnati.

To induce myocarditis, IL-5tg received subcutaneous immunizations on days 0 and 7 of 100 µg myosin heavy chain α (MyHCα) 614 peptide (Ac-SLKL MATL FSTY ASAD; Genscript), or 790 peptide (Ac-IQAQ ARGQ LMRI EFKK)^15^ emulsified in complete Freund’s adjuvant (CFA, Sigma-Aldrich) supplemented to 5 mg/ml heat-killed Mycobacterium tuberculosis strain H37Ra (BD Biosciences). On day 0, mice also received 500 ng pertussis toxin intraperitoneally (List Biologicals)^16^.

Pre-challenge and weekly during the protocol venous blood was collected from mice via submandibular puncture. If complete blood counts were being performed, blood was collected in K_2_EDTA coated tubes (BD Biosciences), while for serum blood was collected in tubes coated with clot accelerator and serum separator gel (BD Biosciences). Samples were inverted in tubes to mix with coating and allowed to settle for 30-60 minutes. The sample was then centrifuged at 1000g for ten minutes at 4°C. Sera were aliquoted to sterile tubes and stored at –80°C until use in assay.

At sacrifice, the hearts were perfused with DPBS (Gibco) supplemented with 0.9mm CaCl_2_(Alfa Aesar), and collected for single cell suspension preparation, histology, and/or RNA.

### Cell free DNA in serum

Sera were warmed to room temperature and diluted 1:20 in assay buffer 30-60 minutes before assay. Quantification of cell free dsDNA performed by fluorometry using PicoGreen assay following the manufacturer’s instructions (Quant-iT PicoGreen dsDNA Kit; Invitrogen). Plates were read on GloMax Multi Detection System (Promega) at wavelengths of 480 nm and 520 nm for excitation and emission, respectively. Fluorescence values were subtracted from sample/standard curve fluorescence values and concentration of cell free DNA was calculated from the standard curve. All steps of the assay were performed at room temperature.

### Troponin

Sera were thawed to room temperature and diluted 1:10 in assay diluent 30 minutes before assay. Quantification of cardiac troponin-I was performed by spectrophotometry using Mouse Cardiac Troponin-I ELISA Kit following the manufacturer’s instructions (CTNI-1-US; Life Diagnostics). Absorbance of wells was measured on a GloMax Multi Detection System (Promega) at wavelength of 450nm. Blank absorbance values were subtracted from sample/standard curve absorbance values and concentration of cardiac troponin-I was calculated from the standard curve. All steps of the assay were performed at room temperature.

### Flow cytometry

Single cell suspension was made from heart as below. Cells were stained with SiglecF-PE (Biolegend), CD19-APC (Invitrogen), CD3-FITC (Invitrogen), Ly6G-BV421 (BD Horizon), CD45-APCcy7 (BD Pharmingen), and 7AAD (Bioscience). Data were collected on Canto3 flow cytometer. Compensation, settings and gating are described in supplementary data (MIFlowCyt format).

### Histology

Tissues were fixed in formalin and embedded into paraffin blocks. Sections were stained with hematoxylin and eosin (H&E) and Trichrome using standard techniques at the Pathology core at Cincinnati Children’s hospital. Anti-MBP immunohistochemistry was performed with antibody gifted by Dr. Elizabeth Jacobsen (Mayo Clinic) using established methods^17^.

### Peripheral blood cell counts

Peripheral blood was collected in EDTA-coated tubes and complete blood counts (absolute white blood cell, red blood cells and platelet count) performed using automated cell counter (Heska). A peripheral blood smear was prepared and stained using Diff Quick (Epredia), and manual differential cells count was performed (since eosinophil count was inaccurate on automated cell counter). The absolute count of individual cell types was calculated from absolute white blood cell count from automated counter and manual differential count.

### Single cell suspensions

The heart single cell suspensions were prepared as per 10X genomics single cell protocol (CG00053 Rev C). Briefly, the heart were cutting into halves using 4-chamber cut, and half a heart was saved for histology, while other half for preparation of single cell suspension. Heart tissue was minced, followed by enzymatic digestion (2.2mg/mL Collagenase IV, Worthington and 1.5mg/mL Dispase II, Life Technologies) at 37°C for 45 minutes. Subsequently, samples were filtered through a 40µm filter, and red blood cells were lysed (eBioscience RBC Lysis Buffer, Fisher Scientific). Cells were then resuspended in RPMI supplemented with 10%FBS and filtered through 30 µm MACS cell strainer (MACS Filters Miltenyi Biotec). The cells were counted to determine concentration and viability using a hemocytometer after which they were subsequently fixed for chromium fixed RNA profiling.

### Fixation of Heart Single Cell Suspension for Chromium Fixed RNA profiling

The fixation was done following 10X genomics Fixation of Cells & Nuclei for Chromium Fixed RNA Profiling protocol (CG000478 | Rev C). Briefly, the heart single cell suspensions were centrifuged at 400g and resuspended in Fixation Buffer, followed by storage at 4°C for 16-24 hours. Following a spin at 850g for 5 minutes at room temperature (22°C), the sample pellet is resuspended in chilled Quenching Buffer, and cell concentration is determined. Pre-warmed Enhancer and glycerol (Thermo scientific) are added prior to storing cells -80°C.

### Workflow for Single-Cell RNA-Seq Data Processing and Analysis

For each mouse, half of the heart was fixed for histology and the other half was used for single cell suspension and fixation in Chromium flex kit and stored at –80°C. Once histological assessment was performed, we selected which mice to subject to single cell RNA sequencing. Library preparation and sequencing were performed at the Single Cell RNAseq Core at Cincinnati Children’s Hospital. The fixed RNA profiling assay was performed according to the manufacturer’s instructions (Chromium Fixed RNA Profiling Reagent Kit User Guide, 10x Genomics). Briefly, individual suspensions of fixed cells were subjected to sample-barcoding (BC001-BC004) using the Chromium Fixed RNA Kit, Mouse Transcriptome (PN-1000496). Mouse whole transcriptome probe pairs were used for overnight probe hybridization. Next, the barcoded samples were pooled, washed, and subjected to gel bead-in-emulsion (GEM) generation using the Chromium Next GEM Single Cell Fixed RNA Sample Preparation Kit (PN-1000414) and the Chromium Next GEM Chip Q Single Cell Kit (PN-1000418 / PN-1000422). The cells were resuspended in a master mix and loaded together with partitioning oil and gel beads into the chip to generate GEMs. Upon the fixed cells and gel beads entering a droplet, the gel beads dissolved, releasing single cell barcoding primers, and the fixed cells lysed, exposing the RNA with probe pairs hybridized to it. The GEMs were collected and incubated in a thermocycler, allowing ligation of the probe pairs followed by hybridization and incorporation of the single cell barcoding primers. The single cell barcoding primers incorporated partial Read 1T, a 16-nucleotide 10x GEM Barcode, a 12-nucleotide unique molecular identifier (UMI), and partial Capture Sequence 1 to the ligated probe pairs. Next, the GEMs were broken, and the cell-barcoded molecules were cleaned up with Silane DynaBeads, then subjected to pre-amplification and SPRIselect reagent size selection. Finally, a gene expression library was constructed. P5, P7, i5 and i7 sample indexes, and Illumina TruSeq Read 1 sequence (Read 1T) and Small Read 2 (Read 2S) sequences were added to generate Illumina sequencer-ready libraries using the Dual Index Kit TS Set A (PN-1000251). The samples were run on one lane of a 10B flow cell on the NovaSeq X Plus sequencer with the following sequencing parameters: R1: 28 cycles, i7: 10 cycles, i5: 10 cycles, R2: 90 cycles.

The Cell Ranger software package from 10x Genomics v.8.0 was utilized to process the raw FASTQ files generated from single-cell RNA sequencing and aligned the sequencing reads to a mouse mm10 reference genome. Additionally, the Cell Ranger performed the initial filtering of low-quality or empty droplets to retain valid cells, as well as filtering genes based on expression levels to focus on those with significant expression that ensured high-quality, filtered data, ready for downstream analysis and interpretation in Seurat.

The Seurat package v5.0.2 was used to preprocess and analyze single-cell RNA-seq data obtained from Cell Ranger. First, the dataset was filtered to retain cells with more than 100 RNA counts and less than 15% mitochondrial gene expression to eliminate potential low-quality cells. The data was normalized, and highly variable features were identified. Subsequently, the data was scaled, and principal component analysis (PCA) was performed. Dimensional reduction to form the uniform manifold approximation and project (UMAP) utilized the top 20 calculated dimensions.

The Doublet Finder package V. 2.0.4 was used to remove doublets^18^ using 8% expected doublet rate formation parameter. Samples were normalized using the sctransform approach with default settings^19^. Dimensional reduction was then performed using the UMAP and utilized the top 30 calculated dimensions and a resolution of 0.4. Data integration was performed in Seurat^20,21^ to merge the single-cell RNA-seq datasets from different conditions, batches, or experiments, facilitating joint analysis. It identifies common features across datasets to align and correct technical differences, enabling the comparison and analysis of cells from disparate sources.

Cell type identification were identified using a combination of canonical markers of cell lineages provided in Table S1, the SingleR (v2.2.0) R package^22^ with correlations of the single-cell expression values with transcriptional profiles from pure cell populations in the Immgen^23^ and the FindAllMarker function in Seurat to confidently annotated each cell cluster.

Differential gene expression utilized the Wilcoxon rank-sum test on count-level mRNA data. For differential comparison across conditions the FindAllMarkers function in the Seurat package was used on of the integrated sctransform-normalized data, employing the log-fold change threshold >0.2, minimum group percentage = 10%, and minimum percentage difference in the fraction of detection between two groups=20%. The differential gene expression analysis was followed by Gene Ontology (GO) enrichment analysis to identify overrepresented biological processes. The analysis was conducted using R with the clusterProfiler(v.4.3.1)^24^, org.Mm.eg.db (v.3.17.0), and Annotation (v.1.64.1). Genes were filtered from the differential expression results to include only those with an average log2 fold change greater than 0.5.

## Results

### Eosinophilic experimental autoimmune myocarditis

In order to mimic hypereosinophilia-associated heart disease, we adapted the EAM model. Hypereosinophilic mice (CD2.IL5 tg) were challenged with a myocarditic (M) and non-myocarditic control (C) peptide from cardiac alpha myosin (**Figure 1A**). Following antigen challenge, mice developed peripheral blood leukocytosis, represented by lymphocytosis, neutrophilia and eosinophilia (comparable between two peptides since both peptides were emulsified in complete Freund adjuvant, CFA, **Figure 1B**). However, at day 21, while mice challenged with control peptide do not show heart inflammation, heart histology in the majority of mice challenged with the myocarditic peptide showed inflammation of all three layers-endocardium, myocardium and epicardium (**Figure 1C**) which was eosinophil-rich (highlighted by anti-MBP staining) and associated morphologically with cardiomyocyte death (**Figure 1C**). From nine experiments performed to date with a total of 45 mice challenged with myocarditic peptide, the average disease penetrance (per cent of mice with heart inflammation by histology) is 56%. In contrast, none of the mice challenged with the control peptide (n=31) developed myocarditis (P<0.0001, Fisher’s exact test). No difference between male and female mice was seen in penetrance or severity of disease. Furthermore, analysis of other organs failed to reveal any destructive inflammation in kidneys, spleen, liver or skeletal muscle (data not shown). In a separate experiment, mice were sacrificed at day 42, and those challenged with myocarditic peptide histologically showed increased fibrosis (**Figure 1C**). Flow cytometry of cardiac single cell suspensions showed increased proportion of hematopoietic (CD45+), T-cells (CD3+) and eosinophils (Siglec-F+) (**Figure 1D**), with no statistically significant difference in neutrophils and B-cells (data not shown). Furthermore, based on the level of surface Siglec-F, eosinophils in spleen show two populations with large Siglec-F low and smaller Siglec-F high (activated population), while eosinophils in the heart were uniformly activated, Siglec-F high (**Figure 1D)**.

**Figure 1.**
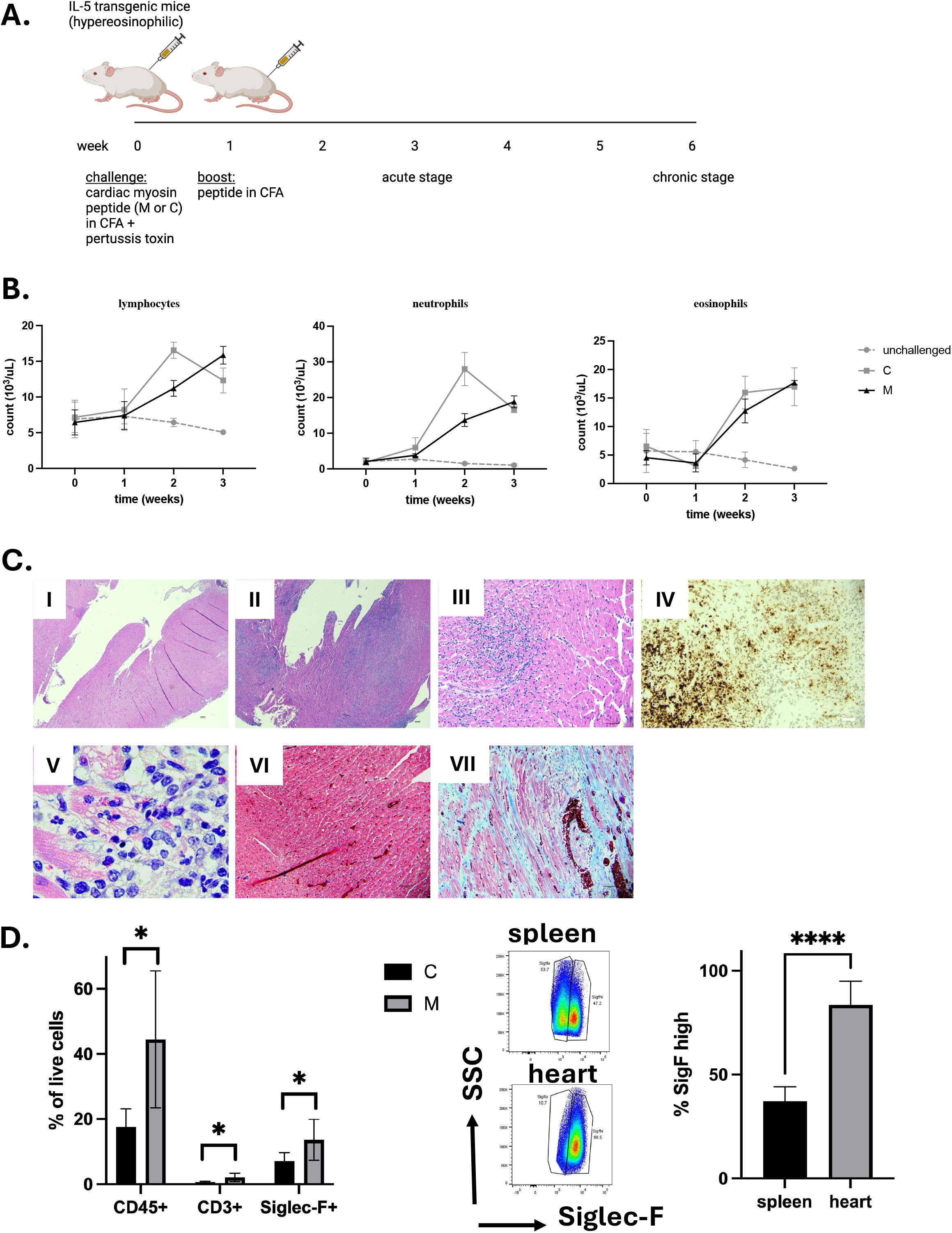
Hypereosinophilia-associated heart disease model. In **A**, schematic representation of the model is shown. M= myocarditic peptide; C= control peptide; CFA= complete Freund adjuvant. In **B**, white blood cells were counted using automated cell counter followed by manual differential on peripheral blood smears from unchallenged (gray dotted line), and mice challenged with control (C, gray solid line) and myocarditic (M, black solid line) peptide. Shown are lymphocytes, neutrophils and eosinophils. Data are average +/-SD from 3 experiments with 2, 13 and 15 mice total (unchallenged, C and M, respectively). In **C**, histologic assessment from the heart is shown: H&E (I-IV) and Trichrome staining (V-VI) of heart tissue 21 (I-IV) and 42 (V-VI) days after challenge with myocarditic (II-IV, VI) and control (I, V) peptide. Magnification: 40x (I-II, scalebar 100 µm), 200x (III, V-VI, scalebar 50 µm), 1000xoil (IV, scalebar 20 µm). Representative mice from 31 and 45 challenged with control and myocarditic peptide are shown. In **D**, heart cells were analyzed by flow cytometry. Single cell suspensions of hearts from mice challenged with control (C) and myocarditic (M) peptide were subjected to flow cytometry for hematopoietic cells (CD45+), T-cells, (CD3+), eosinophils (SiglecF+), B-cells (CD19+, data not shown), neutrophils (Ly6Ghi/SiglecF-, data not shown). In middle panel, the level of Siglec-F is compared in spleen and heart eosinophils in mice challenged with M peptide, and % Siglec-F high cells plotted. ^*^ = P<0.05; ^****^ = P<0.0001. Data are from 6-7 mice per group.

Histologic assessment showed cardiomyocyte dropout, and damaged cardiomyocytes some with clear eosinophil free extracellular granules (e.g. **Figure 1C-v**). Thus, we hypothesized there is tissue (particularly cardiomyocyte) damage. In order to assess for signs of tissue damage, we measured the level of cell free DNA (cfDNA) circulating in peripheral blood (**Figure 2A**). The level of circulating cfDNA did not change significantly over time in unchallenged mice or mice challenged with the C peptide. However, the cfDNA levels increased 22.2 ± 10.2 fold at 1 week, 47.8 ± 11 at 2 weeks and 50.5 ± 9.5 at 3 weeks (P<0.0001 by 2-way ANOVA) in mice challenge with the M peptide. In order to assess for cardiomyocyte damage, we measured the level of cardiac troponin in the serum of mice. The level of troponin was undetectable at baseline, and increased to measurable level at 3 weeks in mice challenged with M peptide (**Figure 2B**). In contrast, unchallenged mice and C-peptide challenged mice did not have measurable troponin. Together, these data indicate that mice challenged with myocardiatogenic peptide experience cardiomyocyte damage.

**Figure 2.**
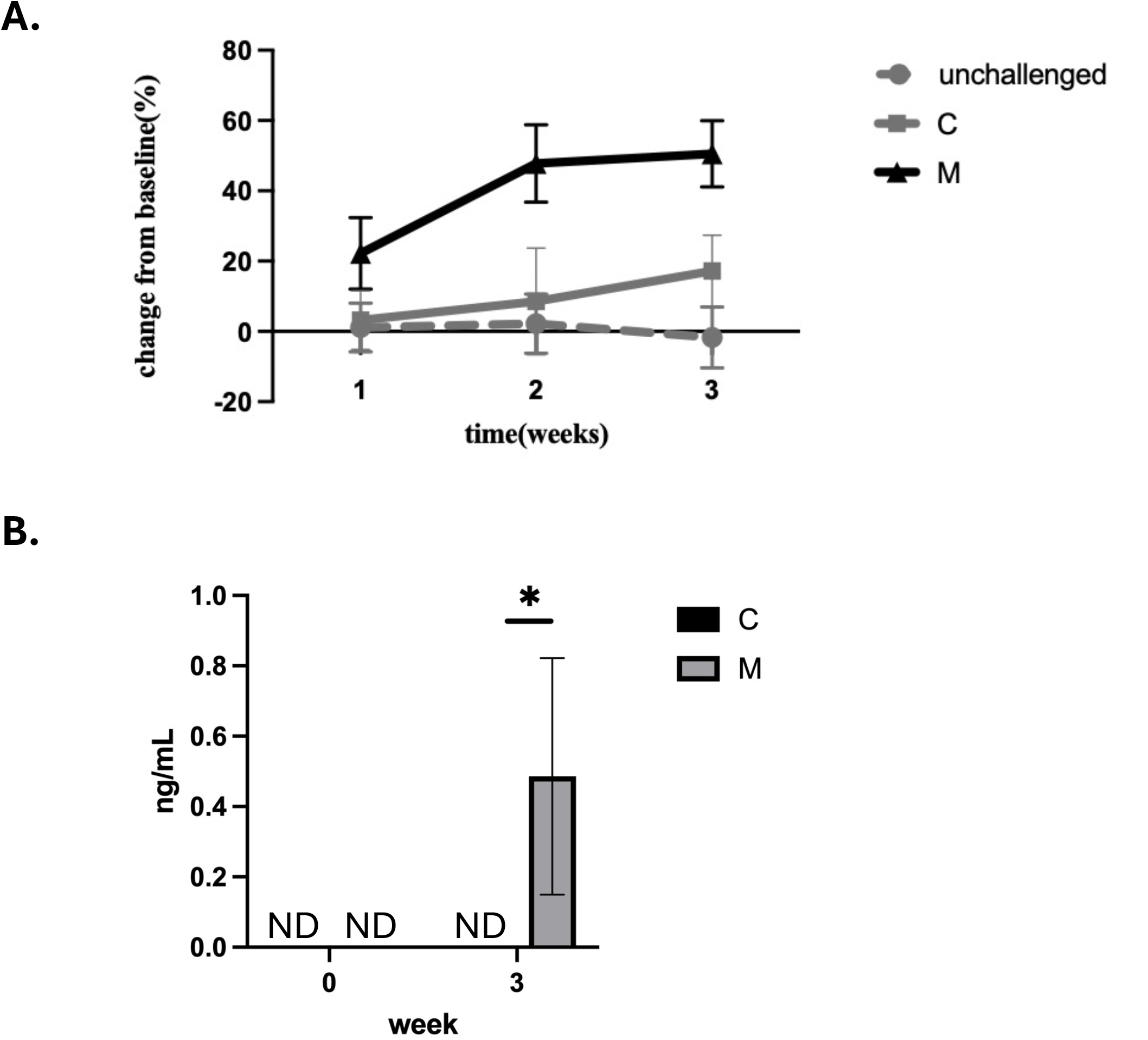
Tissue damage in eoEAM model. In **A**, cfDNA was measured in serum of mice challenged with the M peptide (black solid line), C peptide (gray solid line) or unchallenged mice (gray dotted line) over 3 weeks of challenge. Data are shown for % change over baseline for each mouse, with mean and SD of 3 experiments (with 3-4 mice per group in each experiment). P<0.0001 by 2-way ANOVA. In **B**, troponin was measured in the serum of mice challenged with myocarditic (M) or control (C) peptide at baseline and peak inflammation time point (week 3). Asterisk signifies P<0.05.

In summary, the EHD model shows eosinophil-rich inflammation associated with cardiomyocyte necrosis (early) and dropout with replacement fibrosis (late), and there is troponin detectable in the serum indicating CM damage. Thus, we now have a model to most efficiently and thoroughly test the role of eosinophils in heart inflammation.

### Role for eosinophils in heart inflammation

Effect seen in IL-5 transgenic mice can be due to direct effect of IL-5, or via cells it activates including B-cells and eosinophils. In order to test the role of eosinophils in heart inflammation, we used constitutively eosinophil-deficient mice (ΔdblGATA) challenged with the myocarditic peptide (**Figure 3A**). Significantly fewer ΔdblGATA mice had heart inflammation and those that did had only rare eosinophils (as seen by H&E and MBP staining, **Figure 3B**) and no definitive cardiomyocyte damage. Thus, these data indicate that eosinophils are critical for heart inflammation in hypereosinophilia-associated heart disease.

**Figure 3.**
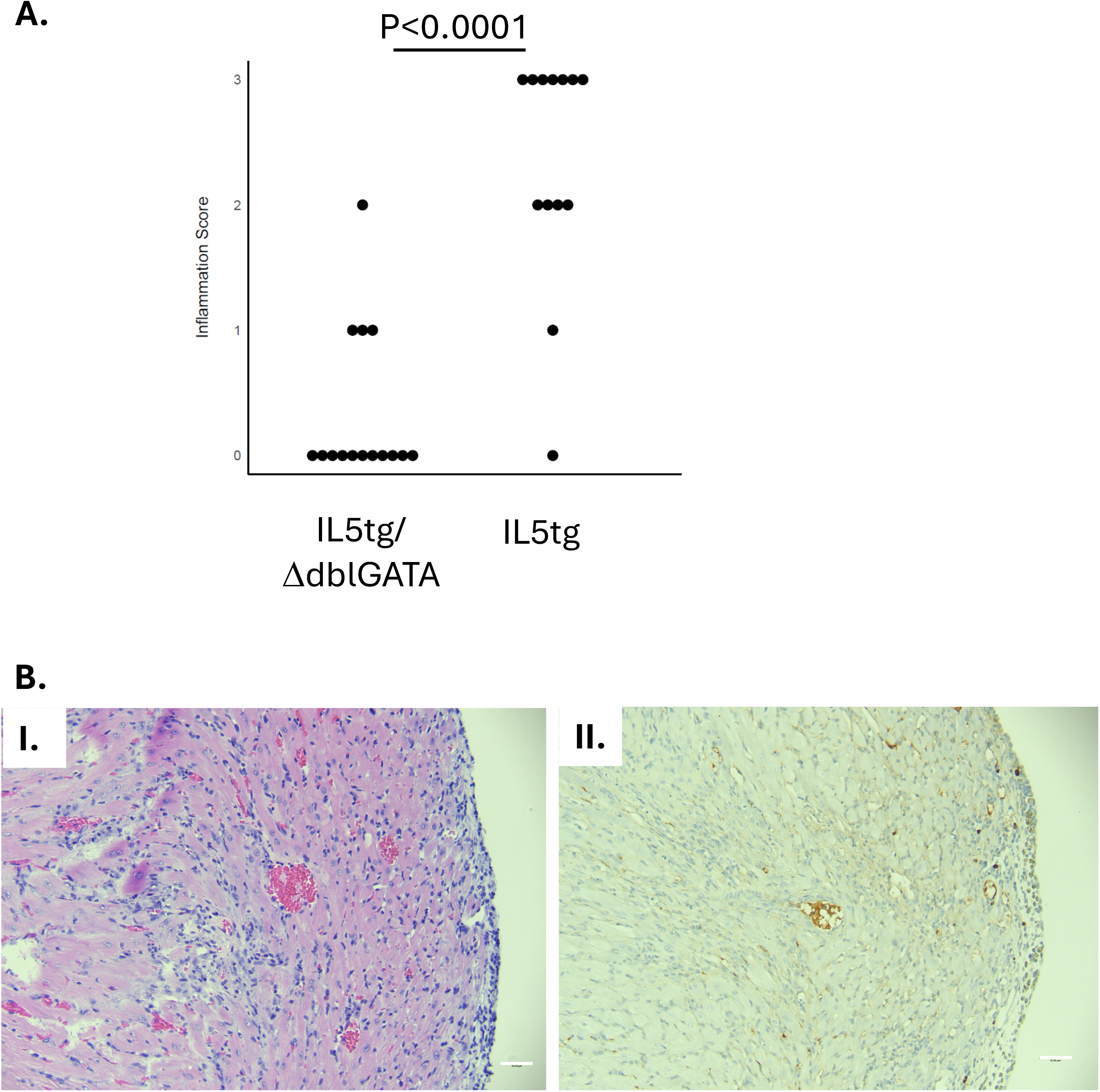
Heart inflammation in ΔdblGATA mice challenged with myocarditic peptide. In **A**, heart inflammation was assessed by histology (semiquantitative assessment by observed blinded to treatment: 0= none, 1= mild, 2= moderate, 3= severe). In **B**, histology of the IL5tg/ΔdblGATA mouse with worse inflammation is shown with H&E (I) and MBP (II) staining. Magnification: 200x; scalebar 50 µm.

### Single cell RNA sequencing

To interrogate cellular composition and diversity in their transcriptome profile, we performed scRNAseq using hearts from mice challenged with control (C) and myocarditic (M) peptide. Single cell suspensions (16,000 cells/mouse) from hearts of challenged mice were loaded into the 10x genomics chromium 3’ expression system, and their libraries were sequenced for downstream analysis. A combined total of 25,885 cells from C (n= 11,770) and M (n= 14,115) hearts were analyzed using the Seurat R package and unbiased clustering yielded 17 clusters (**Figure 4A**). We annotated clusters using a combination of canonical markers, comparison to Immgen database and confirmed with FindAllMarkers function in Seurat. By comparing the relative proportions of each cell type in both groups, we found that mono/mac, neutrophils, T cells and eosinophils are enriched, B-cell are not changed, and fibroblasts and endothelial cells are decreased in myocarditic compared with control hearts (**Figure 4B**). These data suggest hematopoietic cells, especially mono/mac, granulocytes and T-cells, have a significant role in pathogenesis of the myocarditis.

**Figure 4.**
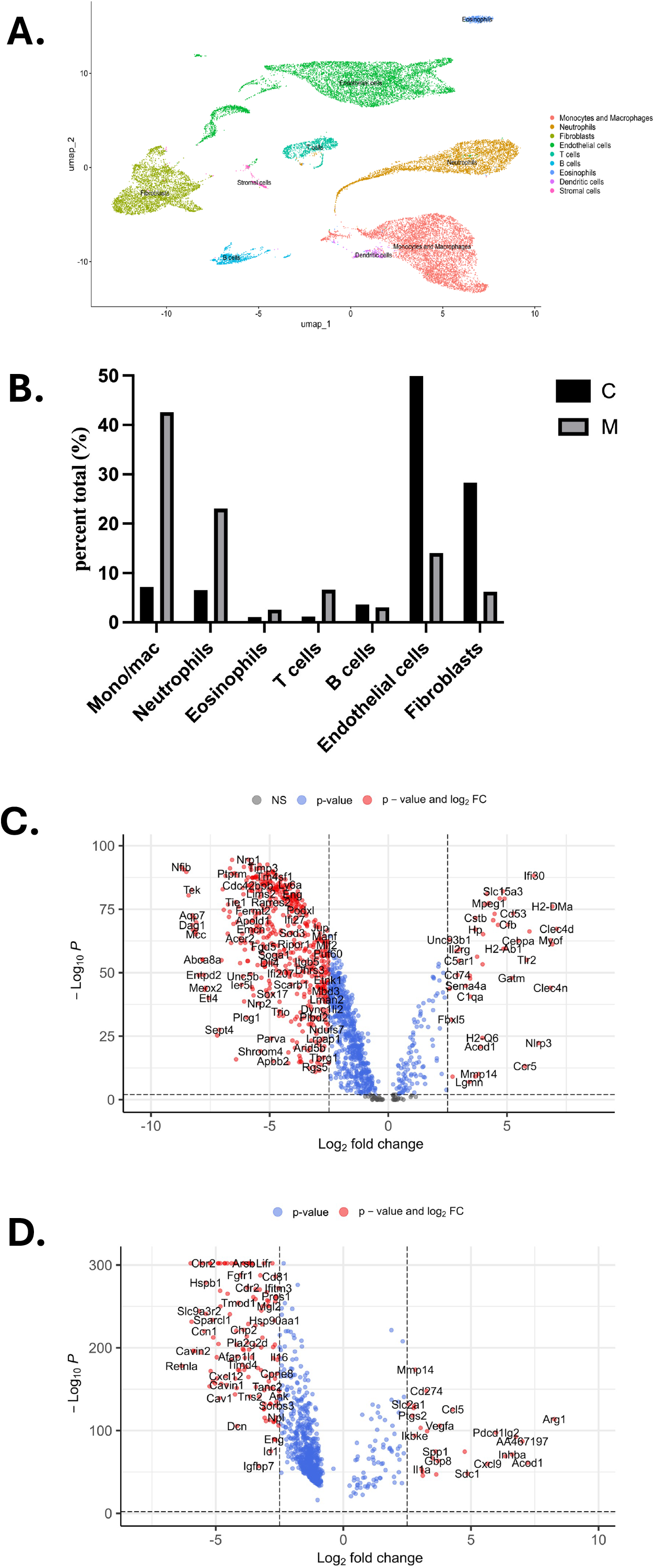
Single cell RNA sequencing. In **A**, cluster analysis of cells from the heart of myocarditic and control peptide challenged mice is shown. In **B**, proportion of cell types in the heart of myocarditic and control peptide challenged mice is shown. In **C** and **D**, volcano plot of differentially expressed genes in eosinophils (C) and monocytes/macrophages (D) is shown.

To determine the functional roles of the enriched cell types, we compared gene expression profiles between the two groups. We first focused on eosinophils because experiments in eosinophil-deficient mice demonstrated their important role in heart inflammation. We found 169 upregulated and 1139 downregulated genes in eosinophils from heart of mice challenged with myocarditic peptide compared with control peptide (**Figure 4C**). There was increased expression of genes associated with complement activation and pathogen-associated molecular pattern recognition. Furthermore, there was increased expression of CD274 (PDL1) consistent with an activated phenotype^25^.

Next we turned to monocytes/macrophages which were the most abundant cells in hearts from mice challenged with myocarditic mice. We found 103 upregulated and 790 downregulated genes in mono/mac from heart of mice challenged with myocarditic peptide compared with control peptide (**Figure 4D**). The top differentially expressed gene was arginase-1, consistent with M2 phenotype^26^. Together, these data show that eosinophils have an activated phenotype and that macrophages are M2 skewed.

## Discussion

In this manuscript, we used a model of hypereosinophilia-associated heart disease to understand the molecular and cellular mechanisms of disease.

As the NIH Taskforce on Research needs of Eosinophil-Associated Diseases (TREAD) and recent RE-TREAD reported, there is a paucity of preclinical models that adequately replicate cardiac disease in hypereosinophilia, and development of these models would enable mechanistic studies aiming to develop targeted therapies^8,9^. Attempts to model hypereosinophilia-associated heart disease in mice include spontaneous eosinophilic myocarditis in DBA/2 (D2) mice ^27^, baseline heart disease in aging IL-5 transgenic mice^28^, and a model of antigen (cardiac myosin)-induced autoimmune EM elicited in IL-5 transgenic mice ^12,29^. The D2 model is limited in that it lacks the endocardial thrombosis, fibrosis, and systemic aspects typical of HES; furthermore, this model develops very early and resolves on its own by ∼3 months of age ^30^. Diny et al have shown that IL-5 transgenic mice develop worsening left ventricular function with age; however, no thrombosis or fibrosis was seen^28^. Studies in myosin-challenged IL-5 transgenic mice developed severe inflammation, followed by dilated cardiomyopathy and fibrosis^12^. It is important to note here that Diny et al used mice where the IL-5 transgene is driven by the CD3 promoter^31^, while our studies use mice where the IL-5 transgene is driven by the CD2 promoter^13^. The level of eosinophilia differs between the two mouse lines, with CD3.IL5tg mice having higher baseline circulating eosinophil levels than CD2.IL5tg (40-60% versus 20-30% range, respectively). Thus, findings from one line cannot be directly assumed to translate to the other line.

Eosinophils have been shown to play both host protective and destructive roles in different models, and the mechanism of their involvement involves multiple effector functions including contributing to antigen presentation and modulation of adaptive immune responses; damage to tissues by cytotoxic granule proteins or antibody-dependent cellular cytotoxicity; and promoting inflammation, thrombosis and/or tissue repair and angiogenesis via secretion of cytokines, chemokines and tissue factor^32,33^. For instance, eosinophils and/or their granule proteins are cytotoxic in variety of scenarios, including data in cardiomyocytes^34,35^. Thus, it was critical to test whether eosinophils are protective or detrimental in hypereosinophilia-associated heart disease. Our studies show that eosinophils are critical for inflammation in that both the occurrence and severity of inflammation are decreased in ΔdblGATA eosinophil-deficient mice.

In order to study the mechanism, we performed scRNAseq analysis from hearts of mice challenged with myocarditic or control peptide. We find increased proportion of hematopoietic cells (mono/mac, T-cells, neutrophils and eosinophils) and decreased proportion of structural cells (fibroblasts and endothelial cells). Assessment of differentially expressed genes identified macrophages as M2-skewed suggesting predominance of type 2 immunity. This is consistent with previous studies showing eosinophils are not just a consequence but also contribute to type 2 immunity^36^.

Furthermore, eosinophils overexpress PDL1 in myocarditic mice, consistent with an activated phenotype^25^. These activated eosinophils have been shown to regulate host defense and immune responses in other disease, and our study now supports their role in heart inflammation.

In summary, eosinophils are required for heart damage in hypereosinophilia-associated heart disease. Additionally, myeloid cells, granulocytes and T-cell cooperatively or independently participate in the pathogenesis of hypereosinophilia-associated heart disease.

## Acknowledgment

The authors thank Drs. Netali Ben-Baruch, Jennifer Felton and Alex Huber for their guidance with single cell sequencing analysis. The project described was supported in part by the National Center for Advancing Translational Sciences (award number 2UL1TR001425, pilot award to N.Z.), National Heart, Lung and Blood Institute (award number HL147898 to N.Z.), National Institute of Diabetes and Digestive and Kidney Diseases (award number P30 DK078392 of the Digestive Diseases Research Core Center in Cincinnati for pathology and flow cytometry cores). The content is solely the responsibility of the authors and does not necessarily represent the official views of the National Institute of Health. Single cell RNA sequencing was performed at the CCHMC Genomics Sequencing Facility (Core Marketplace Research Resource Identifier RRID:SCR_022630). The authors thank Drs Byrd and Hertlein for the use of automated cell counter.

