## supplemental info for "Pathophysiology of hypereosinophilia-associated heart disease"

**Content:**

- Table S1 (marker genes of cell types)
- Supplementary methods: flow cytometry information in MIFlowCyt format

**S Table 1: Marker genes of cell types**

| Cell Type | Marker Genes |
|---|---|
| Eosinophils | Epx, Prg2, Siglecf, Ccr3, Il5ra |
| Neutrophils | Csf3r, S100a8, S100a9 |
| Mast cells | Fcer1a, Mcpt4, Tpsb2, Cpa3 |
| Monocytes & Macrophages | Cd68, Cd168 |
| Dendritic cells | Cd209a, Cd68 |
| B cells | Ighm, Igkc, Jchain |
| T cells | Cd3e, Cd4, Icos, |
| Epithelial cells | Krt14, Krt13 |
| Endothelial cells | Pecam1, Esam, Selp |
| Fibroblasts | Postn, Fbln1, Fbln2 |
| Transitioning fibroblasts | Msn, Lmna, Upk3b |
| Mesothelial cells | Upk3b, Msln, Lrn4 |
| Sensory neurons | Piezo2, Grm5 |

**Minimum Information about a Flow Cytometry Experiment (MIFlowCyt)**

**Eosinophilic experimental autoimmune myocarditis (eoEAM) experiment**

### Introduction

### 1. Experiment Overview

#### 1.1. Purpose

The purpose of the experiment is to immunophenotype and quantify the proportion of infiltrating lymphocytes and myeloid cells by flow cytometric analysis in hypereosinophilia- associated heart disease of eosinophilic experimental autoimmune myocarditis (eoEAM) mouse model. Cardiac and splenic cells were stained with the same appropriate antibodies and similar gating strategies are applied to both tissues.

#### 1.2. Keywords

Heart, spleen, cardiac cell, splenic cell, T cell, B-cell, eosinophil, neutrophil

#### 1.3. Experiment Variables

0.5-1.0x10^6 single cells from spleen or heart

#### 1.4. Organization

1.4.1. Name: University of Cincinnati, College of Medicine, Department of Pathology and Laboratory Medicine

1.4.2. Address: 3230 Eden Ave, Cincinnati, OH 45267; USA

#### 1.5. Primary Contact

1.5.1. Name: Nives Zimmermann

1.5.2.

#### 1.6. Date

| Injection dates performed | 1st injection | 2nd injection | Sacrifices |
|---|---|---|---|
| Experiment 1: | 12/1/2022 | 12/7/2022 | 12/21/2022 |
| Experiment 2: | 5/5/2023 | 5/12/2023 | 5/26/2023 |
| Experiment 3: | 7/16/1013 | 7/13/2023 | 7/27/2023 |

Flow cytometry analysis performed 3 weeks post injection.

#### 1.7. Conclusions

N/A

**1.8. Quality Control Measures**
Splenic cells collected from challenged mice were used in order to provide a staining control such as fluorescence minus one (FMO), and single staining control. Cardiac and splenic cells were used for unstained control.

**1.9. Other Relevant Experiment Information**
N/A

**2. Flow Sample/Specimen Details**

**2.1. Sample/Specimen Material Description**

2.1.1. Biological Samples

*2.1.1.1. Biological Sample Description*: 0.5-1.0x10^6 of cells collected in heparinized in 12x75 FACS tubes for staining

*2.1.1.2. Biological Sample Source Description*: Mus musculus, BALB/c

*2.1.1.3. Biological Sample Source Organism Description:*

- *Taxonomy*: Mus musculus , BALB/c background
- *Age*: >6wk-old mice
- *Gender*: Female and male
- *Phenotype*: white
- *Genotype*: CD2.IL-5 transgenic mice (CD2-IL-5tg)
- *Treatment*:
  - 100 µg myosin heavy chain α (MyHCα) 614 peptide (Ac-SLKL MATL FSTY ASAD; Genscript) for M- group.
  - or 790 peptide (Ac-IQAQ ARGQ LMRI EFKK) for C-group
  - Unchallenged group has no treatment
- *Irradiation*: N/A
- *Other Relevant Biological Sample Source Organism Information:*
  The animal care and house for use at the University of Cincinnati (UC) under Laboratory Animal Medical Services (LAMS).

*2.1.1.4. Other Relevant Biological Sample Information:*
Cage number and mouse ID was used as sample/file identifier.

2.1.2. Environmental Samples
N/A

2.1.3. Other Samples
N/A

**2.2. Sample Characteristics**

Expected/analyzed types of cells: cardiac and splenic cells are stained for lymphoid and myeloid cell characterization

**2.3. Sample Treatment Description**

Single cell suspensions were prepared and the cell concentrations are adjusted to $10^6$/200ul. Fc-blocked using anti-mouse CD16/CD32 antibody (Invitrogen eBioscience) for 10 minutes on ice in FACS buffer. The cells are not washed before the first staining step.

Cells have been incubated for 30 minutes at 4°C in a staining buffer (approx. $10^6$ cells in 200µl of staining buffer). The staining buffer contained a pre-titrated, optimal concentration (≤ 1µg) of a fluorescent monoclonal antibody specific for a receptor or with an immunoglobulin (Ig) isotype-matched control respectively (see below for details).

After the incubation, cells have been washed 1x with 1ml of staining buffer and pelleted by centrifugation (400xg for 5 min); supernatant has been removed.

Finally, cells have been resuspended in 400ul FACS buffer with 7AAD (bioscience ref# 006993-50) and was incubate at RT for 15min prior to flow analysis

**2.4. Fluorescence Reagent Description**

Compensation tube have been created as follows:

| Reporter | PE | APC | FITC | BV421 | PEcy7 | APCcy7 | BV605 | 7AAD | none |
|---|---|---|---|---|---|---|---|---|---|
| Tube#1 | +Ab | - | - | - | - | - | - | - | - |
| Tube#2 | - | +Ab | - | - | - | - | - | - | - |
| Tube#3 | - | - | +Ab | - | - | - | - | - | - |
| Tube#4 | - | - | - | +Ab | - | - | - | - | - |
| Tube#5 | - | - | - | - | +Ab | - | - | - | - |
| Tube#6 | - | - | - | - | - | +Ab | - | - | - |
| Tube#7 | - | - | - | - | - | - | +Ab | - | - |
| Tube#8 | - | - | - | - | - | - | - | +Ab | - |
| Tube#9 | - | - | - | - | - | - | - | - | - |

Each sample has been stained as follows:

| Reporter | PE | APC | FITC | BV421 | PEcy7 | APCcy7 | BV605 | 7AAD |
|---|---|---|---|---|---|---|---|---|
| Sample | +Ab | +Ab | +Ab | +Ab | +Ab | +Ab | +Ab | +Ab |

| Analyte | reporter | Detector | target | Manufacturer | Cat# |
|---|---|---|---|---|---|
| Siglec-F | PE | CD170 (Siglec F) Monoclonal Antibody | eosinophil | Invitrogen | |
| CD19 | APC | Anti-Mouse CD19 antibody | B-lymphocyte | Invitrogen | Cat.#17019382 |
| CD3 | FITC | Anti-Mouse CD3e antibody | T-lymphocyte | Invitrogen | Cat.#11003182 |
| Ly6G | BV421 | Rat Anti-Mouse Ly-6G antibody | Neutrophil | BD Horizon | Cat.# 562737 |
| F4/80 | PEcy7 | anti-mouse F4/80 Antibody | Mono/macrophage | Biolegend | Cat# 123113 |
| viability | 7AAD | | Live/dead | eBioscience | ref# 006993-50 |

| | | | | | |
|---|---|---|---|---|---|
| CD45 | APCcy7 | Rat Anti-Mouse CD45 antibody | Hematopoietic cells | BD Pharmingen | Cat.#1046748 |
| CD49b | BV605 | Anti-Mouse CD49b (Integrin α2) | NK cells | BD | Cat#. 569508 |
| CD122 | BV605 | Rat anti-Mouse | NK cells | BD Biosciences | Cat#BDB745171 |

3. **Instrument Details**

**3.1. Instrument Manufacturer**

Beckton Dickinson

**3.2. Instrument Model**

BD/FACSCanto

Technical specification at: https://www.cincinnatichildrens.org/research/cores/flow-cytometry

**3.3. Instrument Configuration and Settings**

3.3.1. Flow Cell and Fluidics

The instrument has not been altered

3.3.2. Light Sources

The instrument has not been altered

| | | **Number of Fluorescence PMTs** | | | | |
|---|---|---|---|---|---|---|
| **Instrument Name*** | **Instrument Make/Model**** | **UV (355nm)** | **Violet (405nm)** | **Blue (488nm)** | **Yellow-Green (561nm)** | **Red (635nm)** |
| **Canto 3** | BD/FACSCanto | | 3 | 5 | | 3 |

3.3.3. Excitation Optics Configuration

The instrument has not been altered.

3.3.4. Optical Filters

The instrument has not been altered, all filters are original and came with the instrument.

3.3.5. Optical Detectors

The instrument has not been altered. Detector voltages have been set to according to the configuration figure below.

3.3.6. Optical Paths

The instrument has not been altered. The following figure shows the filter and detector configuration:

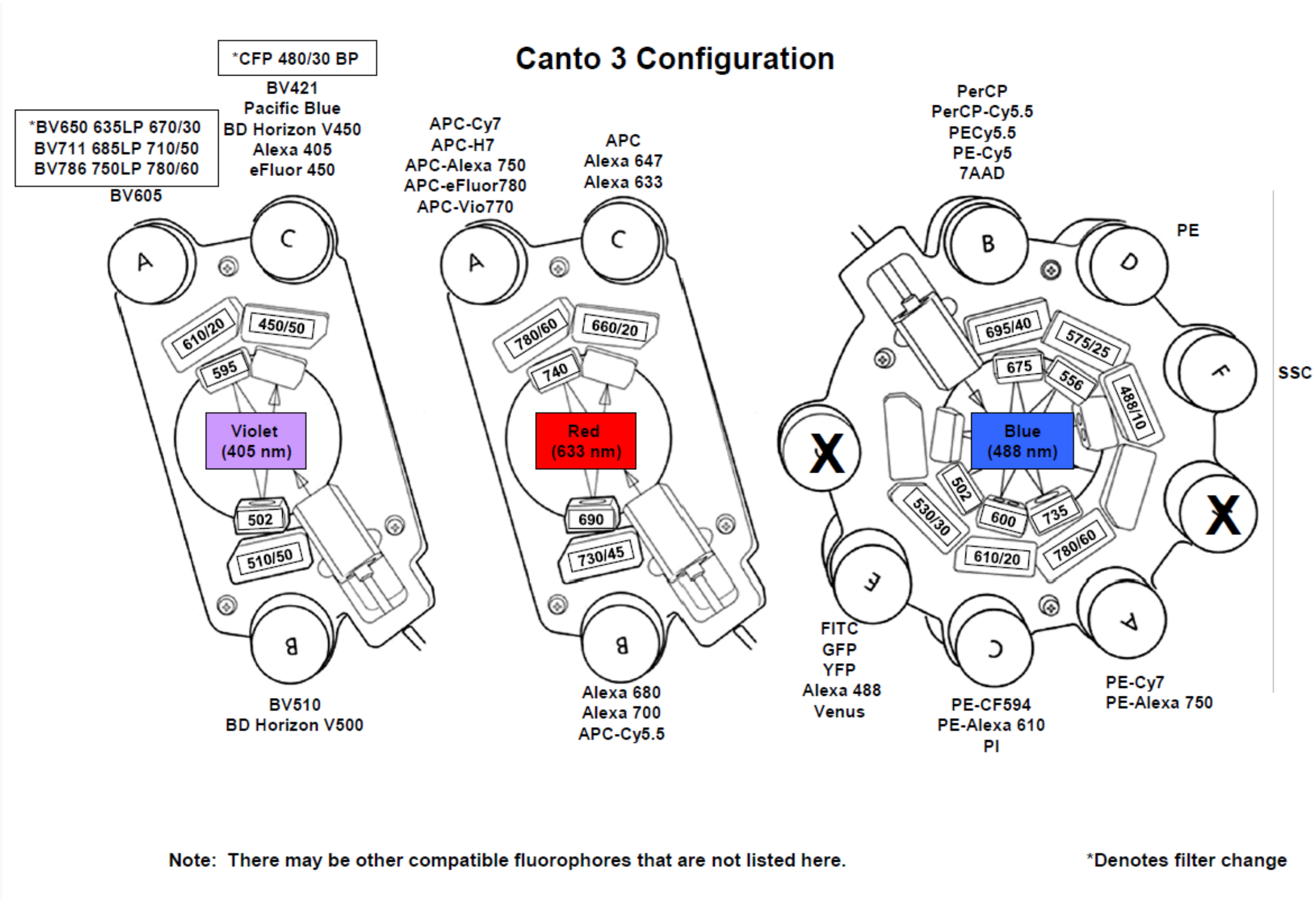


*Figure 1: Optical detectors configuration*

| Detector Array (laser) | PMT | LP Mirror | BP filter | Dye/detector |
|---|---|---|---|---|
| Violet (405nm) | A | 595 | 610/20 | 635LP 670/30<br>BV711 685LP 710/50<br>BV786 750LP 780/60<br>BV605 |
| | B | 502 | 510/50 | BV510<br>BD Horizon V500 |
| | C | - | 450/50 | BV421<br>Pacific Blue<br>BD Horizon V450<br>Alexa 405<br>eFluor 450<br>*CFP 480/30 BP |
| Red (633 nm) | A | 740 | 780/60 | APC-Cy7<br>APC-H7<br>APC-Alexa 750<br>APC-eFluor780<br>APC-Vio770 |
| | B | 690 | 730/45 | Alexa 680<br>Alexa 700<br>APC-Cy5.5 |
| | C | - | 660/20 | APC<br>Alexa 647<br>Alexa 633 |

| | | | | |
|---|---|---|---|---|
| Blue (488nm) | A | 735 | 780/60 | PE-Cy7<br>PE-Alexa 750 |
| | B | 675 | 695/40 | PerCP<br>PerCP-Cy5.5<br>PECy5.5<br>PE-Cy5<br>7AAD |
| | C | 600 | 610/20 | PE-CF594<br>PE-Alexa 610<br>PI |
| | D | 556 | 575/25 | PE |
| | E | 502 | 530/30 | FITC<br>GFP<br>YFP<br>Alexa 488<br>Venus |
| | F | - | 488/10 | |

#### 3.4. Other Relevant Instrument Details

https://www.cincinnatichildrens.org/research/cores/flow-cytometry/analyzer
https://www.cincinnatichildrens.org/research/cores/flow-cytometry/software

### 4. Data Analysis Details

#### 4.1. List-mode Data Files

FCS data files can be obtained by contacting Dr. Nives Zimmermann after this work has been published.

#### 4.2. Compensation Description

Compensation has been performed computationally post acquisition according to the following spillover matrix (values in %):

| Show All | 7-AAD-A | APC-A | APC-Cy7-A | BV421-A | BV605-A | FITC-A | PE-A | PE-Cy7-A |
|---|---|---|---|---|---|---|---|---|
| 7-AAD-A | 100 | 7 | 2.5 | 30 | 4.5 | 4 | 4 | 50 |
| APC-A | 0.7699 | 100 | 0 | 0.0513 | 0.1026 | -0.154 | 0 | 0 |
| APC-Cy7-A | 0.546 | 28 | 100 | 0.2427 | 0.546 | 0.1214 | 0.182 | 4.6175 |
| BV421-A | 0.0023 | 0.0537 | 0 | 100 | 0.1331 | 0.1356 | 0.0163 | 0 |
| BV605-A | 8.1811 | 1.1307 | 0 | 20 | 100 | -0.5384 | 2.8559 | 0.8614 |
| FITC-A | 1.5827 | 0.1356 | 0 | 0 | 0.1808 | 100 | 14.9168 | 0.1356 |
| PE-A | 11.7372 | 0.4766 | 0 | 1.0732 | 1.7658 | 2.0315 | 100 | 0.7626 |
| PE-Cy7-A | 0.3514 | 0.0836 | 3.7423 | 0.2175 | 0.2007 | 0.2594 | 1.4336 | 100 |

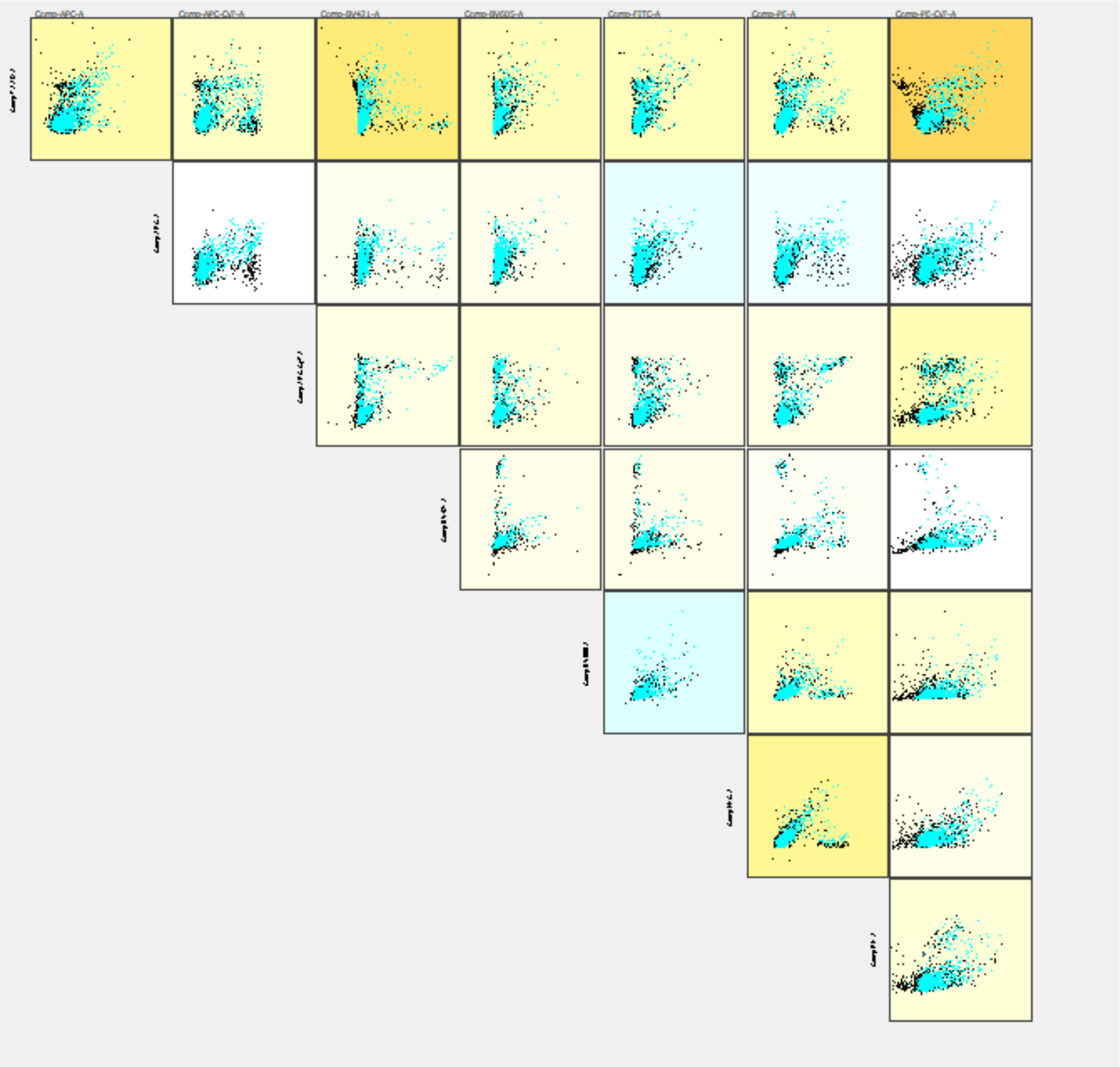


### 4.3. Data Transformation Details

4.3.1.Purpose of Data Transformation
The data has not been transformed
4.3.3.Other Relevant Data Transformation Details
N/A

### 4.4. Gating (Data Filtering) Details

The same gating strategy has been used for all data files (all challenges and unchallenged mice at indicated time point). For simplicity and clarity, we provide details for only one sample gating of a single time point, we include these as images within this document.

4.4.1. Gate Description
The gating strategy involves the following gates:

- Total cells were gathered by FSC-SSC gating; this was called cardiac cells

- From the cardiac cell, single cells were selected by applying FSC-A/FSC-H gating
- From single cells dead cells were removed, and the remaining cells are called viable or Live cells
- From live cells, CD45 positive hematopoietic cells were selected.
- The other cell targets are set as a proportion of CD45 positive live cells
- Activate eosinophil are a proportion of Siglec-F positive cells

4.4.2. Gate Statistics

The following table shows an example of percentages of each of the subpopulations defined by described gates.

| ID# 3Wks T1 | FSC-SSC | FSC-FSC | viable | CD45 | Bcells | Tcells | SiglecF | ly6G |
|---|---|---|---|---|---|---|---|---|
| **total cardiac cells %** | 93.8 | | | | | | | |
| **single cell%** | | 94.4 | | | | | | |
| **Viable%** | | | 84.9 | | | | | |
| **CD45%** | | | | 37.3 | | | | |
| **B-cells%** | | | | | 2.68 | | | |
| **T-cells%** | | | | | | 2.77 | | |
| **Siglec-F%** | | | | | | | 37.3 | |
| **ly6G%** | | | | | | | | 26.5 |

Gate Statistics applies to all used data files in a real experiment description. we only provide a single data file in order to keep this document as a clear and simple example

4.4.3.Gate Boundaries

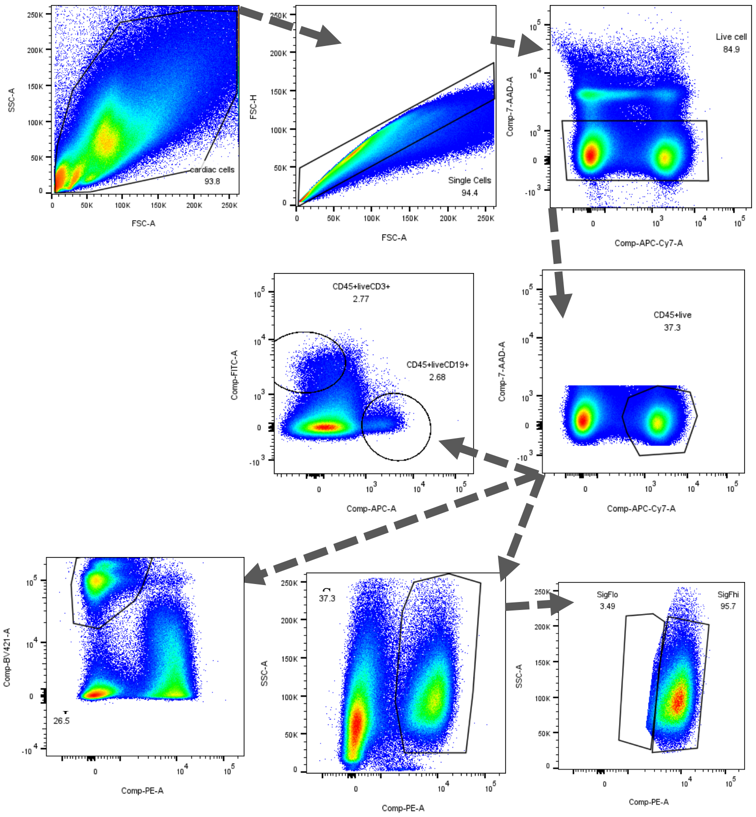

4.4.4.Other Relevant Gate Information

FlowJo workspace files could be obtained by contacting Dr. Nives Zimmermann after this work has been published.
